# Genome-wide discovery of somatic coding and regulatory variants in Diffuse Large B-cell Lymphoma

**DOI:** 10.1101/225870

**Authors:** Sarah Arthur, Aixiang Jiang, Bruno M. Grande, Miguel Alcaide, Anja Mottok, Daisuke Ennishi, Christopher Rushton, Selin Jessa, Prince Kumar Lat, Prasath Pararajalingam, Barbara Meissner, Merrill Boyle, Lauren Chong, Daniel Lai, Pedro Farinha, Graham W. Slack, Jordan Davidson, Kevin R. Bushell, Sohrab Shah, Dipankar Sen, Steven J.M. Jones, Andrew J. Mungall, Randy D. Gascoyne, Marco A. Marra, Christian Steidl, Joseph M. Connors, David W. Scott, Ryan D. Morin

**Affiliations:** Department of Molecular Biology and Biochemistry, Simon Fraser University, Burnaby, BC, Canada.; Centre for Lymphoid Cancer, BC Cancer Agency, Vancouver, BC, Canada.; Genome Sciences Centre, BC Cancer Agency, Vancouver, BC, Canada.; Molecular Oncology, BC Cancer Agency, Vancouver, BC, Canada.

## Abstract

Diffuse large B-cell lymphoma (DLBCL) is an aggressive cancer originating from mature B-cells. Many known driver mutations are over-represented in one of its two molecular subgroups, knowledge of which has aided in the development of therapeutics that target these features. The heterogeneity of DLBCL determined through prior genomic analysis suggests an incomplete understanding of its molecular aetiology, with a limited diversity of genetic events having thus far been attributed to the activated B-cell (ABC) subgroup. Through an integrative genomic analysis we uncovered genes and non-coding loci that are commonly mutated in DLBCL including putative regulatory sequences. We implicate recurrent mutations in the 3’UTR of *NFKBIZ* as a novel mechanism of oncogene deregulation and found small amplifications associated with over-expression of FC-γ receptor genes. These results inform on mechanisms of NF-*κ*B pathway activation in ABC DLBCL and may reveal a high-risk population of patients that might not benefit from standard therapeutics.

## Introduction

It has been established that DLBCL, although genetically heterogeneous, can be robustly divided at the gene expression level into two subgroups based on markers of B-cell differentiation and NF-*κ*B activity pathways, the latter being particularly active in ABC cases^1^. *EZH2^2^, SGK1, GNA13* and *MEF2B*^2^ exemplify genes with mutations restricted to GCB cases, whereas *MYD88*^3^, *CD79B*^4^ and *CARD11*^5^ have been reported as more commonly mutated in ABC. Some DLBCL cases have few (if any) genetic alterations strongly associated with either subgroup, suggesting the possibility of additional genetic or epigenetic changes that shape the malignancy. Similarly, the over-expression of proteins with potential therapeutic and clinical relevance cannot always be explained by known genetic alterations^6^. Gaining a more complete understanding of the genetic features of DLBCL in general and each subgroup in particular should lead to improved methods for this sub-classification and further inform on the molecular and genetic underpinnings of the lymphoma found in individual patients. Such enhancements have the potential to facilitate the development of targeted therapies, such as small molecule inhibitors^7^, new monoclonal antibodies and immunotherapies that target somatic mutations or cell surface proteins^8^.

Although there have now been more than 1000 tumours analysed using targeted strategies such as array-based copy number analysis^9^ or whole exome sequencing (WES)^10^, a limited number of DLBCL genomes have been described to date^11-13^, leaving the potential to uncover new somatic structural variations (SVs), copy number alterations (CNAs) and other *cis*-acting regulatory mutations that may be cryptic to other assays. The search for driver mutations has been further confounded by aberrant somatic hypermutation (aSHM) affecting a substantial number of genes in DLBCL^14^. Specifically, in DLBCL and several other lymphoid cancers the AID enzyme (encoded by *AICDA),* in cooperation with POL*η*, induces mutations in actively transcribed genes with a concentration in the first 1.5-2 kb^15^. Though the repertoire of known aSHM targets in lymphoma continues to grow, it has become apparent that this process can also impact non-genic loci associated with super-enhancers and some aSHM-mediated mutations may have regulatory functions^16,17^.

Whole genome sequencing (WGS) offers the possibility of cataloguing the sites affected by this process along with concomitant determination of genes with potential cis-regulatory effects. We analysed WGS data from 153 DLBCL tumour/normal pairs alongside existing WES data from 191 additional cases to uncover novel driver genes affected by somatic single nucleotide variants or indels, collectively referred to as simple somatic mutations (SSMs). These affected many of the genes that have been ascribed to DLBCL along with 4,386 regions we identified as enriched for somatic mutations, the majority impacting non-coding loci. Analysis of matched RNA-seq data uncovered recurrent structural alterations and mutated loci with potential roles in mediating the transcriptional or post-transcriptional regulation of numerous genes with relevance to DLBCL.

### Integrative analysis of Structural Variation, Copy Number Alterations and Gene Expression

The landscape of somatic CNAs in DLBCL has been addressed by multiple groups^9,18,19^ but owing to the technologies typically used, the breakpoints that underlie these events and putative copy-neutral alterations and smaller focal gains and losses can be missed by array-based approaches^11^. The 153 genomes were analysed for SVs, revealing a total of 13,643 breakpoints (range: 0-390; median 66). We determined the SVs likely to affect specific genes based on their proximity to individual genes (Table 1). As expected, the genes with proximal SVs in the highest number of cases were oncogenes relevant to DLBCL including *BCL2, BCL6, FOXP1* and *MYC.* Tumour suppressor genes (TSGs) typically exhibited focal deletions and commonly exhibited SV breakpoints within the gene body, including *TP53, CDKN2A,* and *CD58* (Extended Data Figure 1).

**Table 1:**
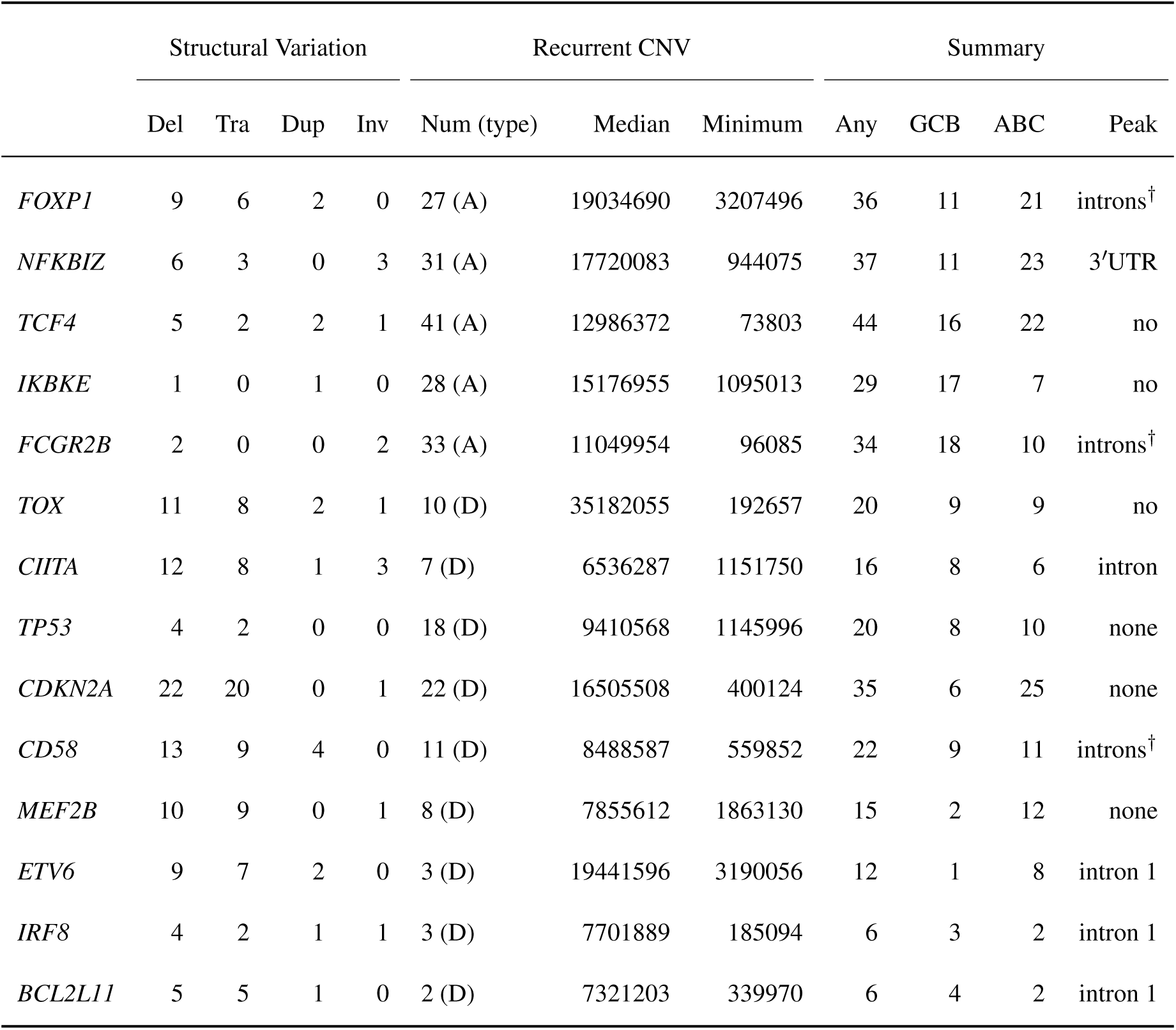
Overview of SVs and CNVs proximal to genes. SVs are separately counted by the type of event as determined by read pairing information. The total number of CNVs in the direction associated with the recurrent alteration (A or D) and the median and minimum of these is shown to highlight the focal nature of some of these events. Tra: translocation; Del: deletion; Dup: duplication; Inv: inversion; A: copy number amplification or gain; D: copy number deletion. †Gene has a visible enrichment of mutations in this region that was not detected by the wavelet approach.

By intersecting with regions affected by recurrent copy number losses, we searched for putative TSGs that might be disrupted by either deletion or SV breakpoints (Table 1). Some of these were separately identified through subsequent analyses (below) revealing patterns of non-silent exonic mutations and/or peaks of non-coding mutations whereas others rarely harboured simple somatic mutations (SSMs) such as SNVs and indels. Many genes impacted by aSHM were also enriched for somatic breakpoints. In contrast, *TOX* and *WWOX* harboured a substantial number of distinct SV breakpoints and several examples of highly focal deletions but rarely harboured SSMs (Figure 1 and Extended Data Figure 1). TOX and WWOX SVs were rare overall, indicating these genes may act as tumour suppressor genes in some DLBCLs. *MEF2B,* a gene that has multiple known mutation hot spots, particularly in GCB DLBCL, also contained several examples of focal deletions or complex SVs. As the function of *MEF2B* mutation in DLBCL has not been fully elucidated^20,21^, this observation strengthens the evidence of its role as a tumour suppressor with its recurrent mutations having a dominant negative effect, but this does not eliminate the possibility of shortened isoforms with an enhanced or distinct activity. Further complicating the matter, *MEF2B* SVs were predominantly found in the genomes of ABC DLBCLs whereas hot spot mutations are a feature of GCB.

**Figure 1:**
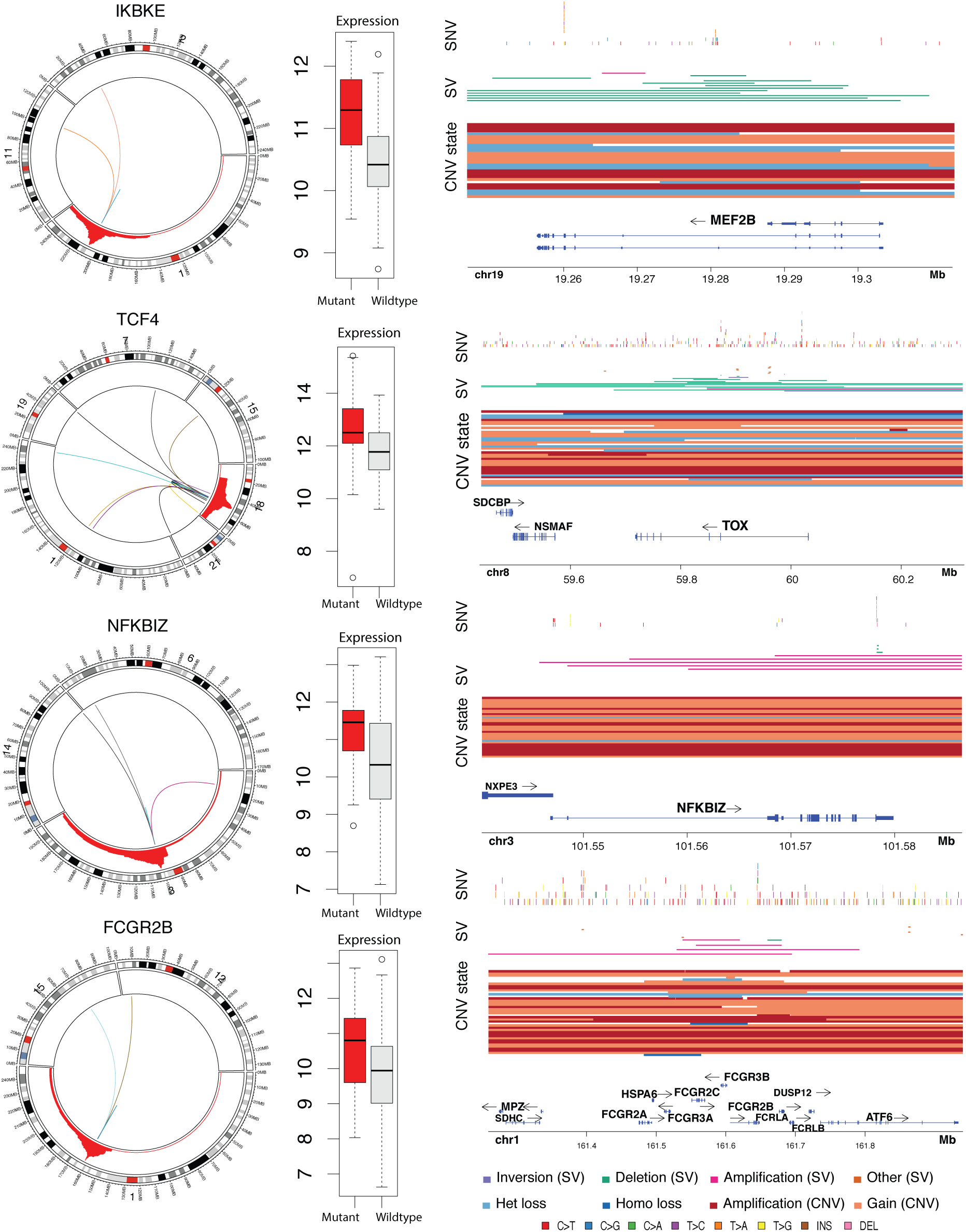
Structural and copy number alterations indicative of oncogenes in DLBCL genomes. Gains and SV breakpoints affecting candidate oncogenes are summarised (left). Only chromosomes involved in at least one SV are displayed for each gene. The red region represents the cumulative number of gains/amplifications encompassing each locus across the cohort of genomes. The expression level of the gene with (red) or without (grey) proximal SV or CNV alterations is shown (centre). Some of the SVs affecting *NFKBIZ* and *TCF4* occur in the gene body and may partially disrupt or alter their normal function. Four examples are shown illustrating the varying patterns of mutation recurrence affecting known and putative DLBCL-related genes (right). *MEF2B* has two main mutation hotspots. This locus and *TOX* are both affected by multiple focal deletions across the cohort of genomes, whereas amplifications and gains of these loci are rare, strengthening their role as tumour suppressor genes. In contrast, *NFKBIZ* showed a striking number of small deletions affecting the 3′UTR. This region of the UTR is also enriched for SSMs. The locus containing *NFKBIZ* is commonly amplified in the genomes and a larger cohort of DLBCLs. The FC-*γ* receptor locus harbours five paralogs. Numerous examples of focal CNVs and a few translocations involving this region were observed. Copy number polymorphisms comprising different combinations of these genes are common in the human population but have been poorly characterised. Paired tumour/normal copy number analysis and SV detection indicates that focal somatic changes in copy number are also common in DLBCL. Despite a preponderance of amplifications, there also appears to be a preference towards focal deletions affecting FC-*γ* receptor genes other than *FCGR2B*.

We next searched for concomitant signals of recurrent copy number gain and SVs proximal to genes, restricting our analysis to regions identified as peaks for amplification by GISTIC in a separate large DLBCL cohort (Ennishi *et al,* unpublished). We utilised RNA-seq-derived expression values from a subset of the cases to infer *cis* effects of these events on expression. This uncovered several genes reported to act as oncogenes through focal amplification, with *IKBKE*, *NFKBIZ*, *FCGR2A/FCGR2B* representing the strongest candidates due to significantly elevated expression in cases having either a gain or proximal SV (Figure 1). This also revealed additional known targets of aSHM (Extended Data Figure 2). Some breakpoints were within the gene body, an observation seen in some known oncogenes such as *FOXP1*^22^. Such events can lead to novel isoforms or fusion transcripts, such as those involving *TBL1XR1*^23^.

The most striking collection of focal gains was those affecting the FC*γ* receptor locus, a complex region of the genome comprising multiple paralogs that have arisen through a series of segmental duplications^24^(Figure 2A-B). These focal gains and, less commonly, deletion events were corroborated by read pairing information in many cases. This observation could be confounded by the presence of germline copy number alterations in this region, thus many of the single copy gains could represent germline events. Using a custom multiplex droplet digital PCR (ddPCR) assay, we confirmed the CNAs and identified four additional examples of amplifications and several additional gains not detected by SNP arrays. Amplifications, but not gains, were significantly enriched among GCB cases and had a striking correlation with elevated *FCGR2B* expression (Figure 2C). Although the prevalence of this genetic alteration is low, we found a compelling trend towards inferior outcome in GCB cases with *FCGR2B* amplification. Taking into account the apparent effect of gains on *FCGR2B* over-expression and deletions on reduced *FCGR2C,* we hypothesised that the tumours benefited from a relative increase in FC**γ**RIIB protein relative to FC**γ**RIIC and used the log-ratio of the expression of these two genes to stratify GCB patients (Methods). This simple classifier showed a strong separation of patients on disease-specific survival that was significant as a continuous variable in a Cox model and showed a strong relationship in univariate Kaplan-Meier analysis across a range of thresholds (Figure 2E-F and Extended Data Figure 3).

**Figure 2:**
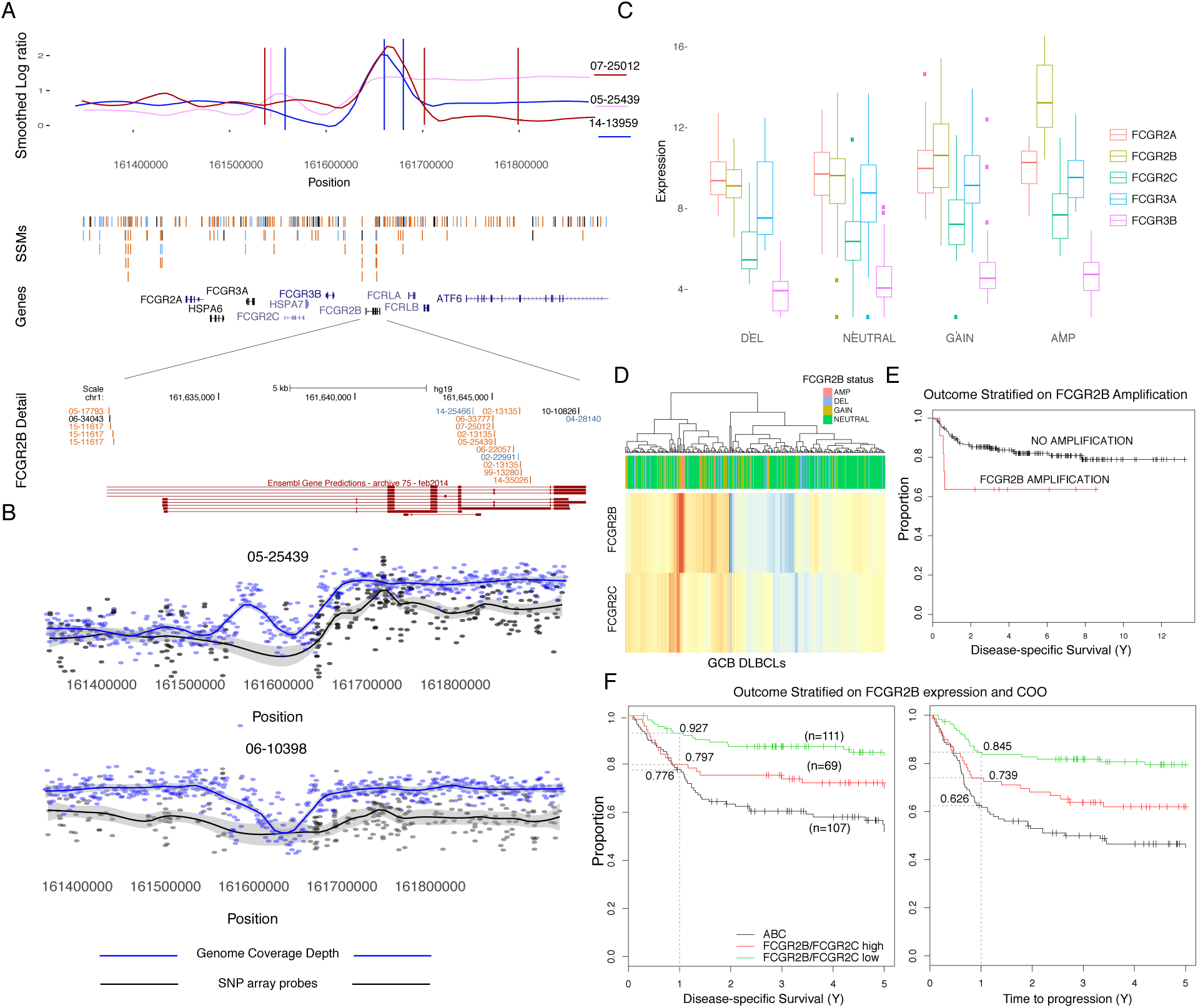
Focal somatic CNAs, SVs and SSMs affecting the Fc*γ* receptor locus. (A) Three genomes with either a focal gain or complex combination of SVs are shown with the smoothed log-ratio of tumour to matched normal read depth confirming they are somatic. Corroborating mate pairs further supported these events, with up to three separate breakpoints detected in one case. The pattern of SSMs in two large introns is shown below, with some of these SSMs predicted to promote intron retention and a shorter CDS, which is annotated by Ensembl (red). (B) Though previous studies have noted some focal gains affecting this region^9^, we note that SNP arrays have poor coverage of this locus. The signal from copy-number probes on SNP6.0 arrays (black) is compared to binned read coverage (blue) for two cases. Due to a lack of constitutional DNA for the validation cohort, we are unable to determine the proportion of single-copy gains and losses that can be attributed to common germline CNVs. We can, however, identify cases with gains exceeding the complement possible through germline CNVs. (C) The local copy number was determined in the validation cohort using ddPCR and amplifications were carefully discriminated from single- or two-copy gains. The expression of each FC*γ* receptor gene is shown with the cases separated by copy number state. Within GCB cases, those with amplifications or gains had significantly elevated *FCGR2B* expression whereas deleted cases showed a trend towards reduced *FCGR2C* expression. (D) Clustering on *FCGR2B* and *FCGR2C* groups amplified cases alongside tumours with gains or no alteration detected, indicating that *FCGR2B* expression may be altered by other avenues. (E) Patients treated with R-CHOP showed an insignificant trend towards inferior outcome. (F) The log-ratio of expression of these two genes was significantly associated with outcome with the *FCGR2B*/FCGR2C-high cases having a shorter DSS and TTP. When the first year is considered, the DSS of high-*FCGR2B* GCB cases was similar to ABC cases.

### Local Mutation Density and Somatic *Cis*-regulatory Variation

We next sought genes with patterns of non-silent mutations, beginning with a meta-analysis of the genomes and all available exome data for recurrently mutated genes. The genes significantly affected by SSMs had mostly been identified in prior studies and a large exome study published while this manuscript was being completed ^10^(Extended Data: Figure 4, Table 1). Among the genes that have limited prior evidence for relevance in DLBCL, many have been implicated in other B-cell lymphomas arising within the germinal centre. For example, *DDX3X, ARID1A* and *HVCN1* which have been reported as recurrently mutated in Burkitt lymphoma (BL)^25^ and follicular lymphoma (FL)^26^.

**Figure 4:**
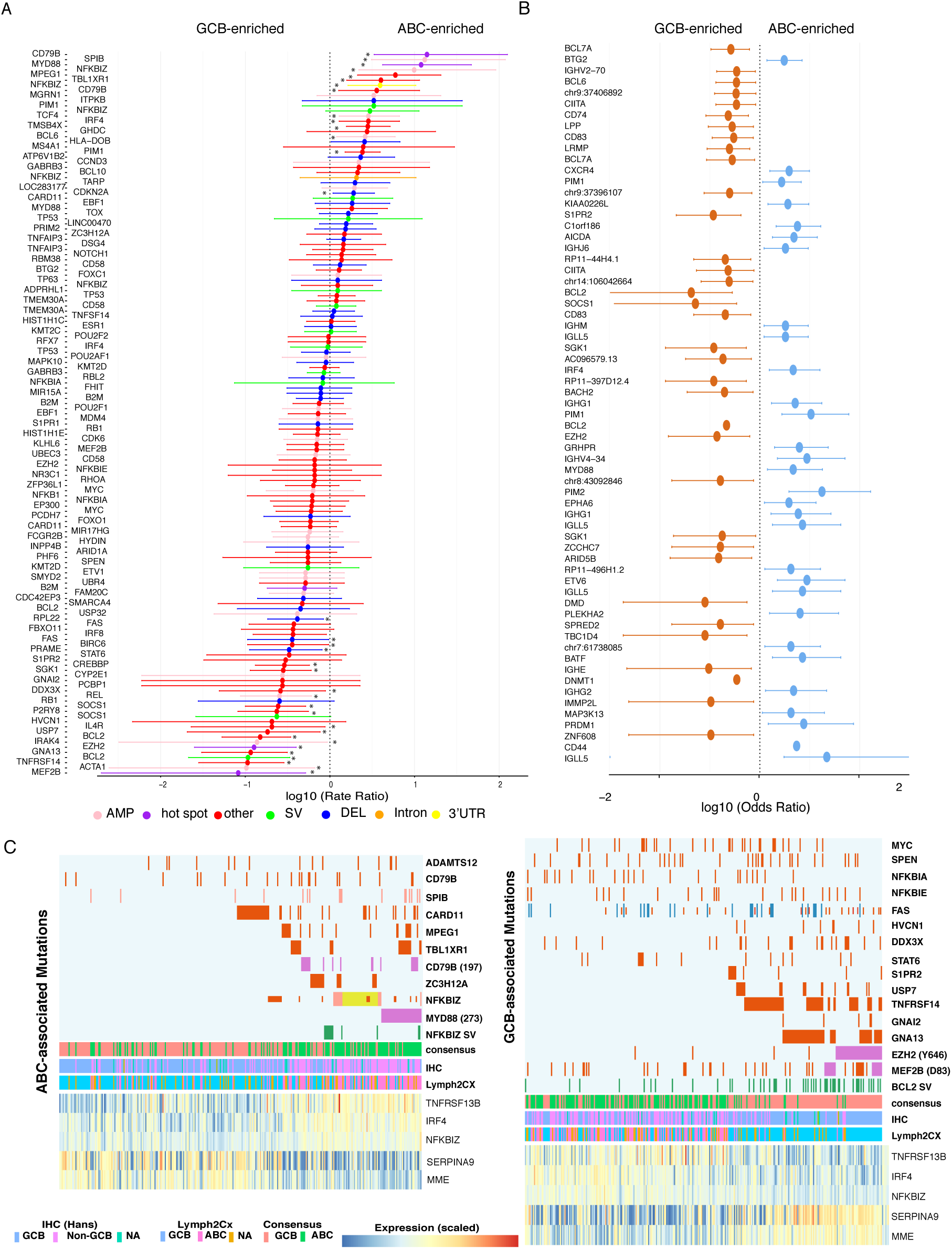
Differences in mutational representation between DLBCL molecular subgroups. (A) Non-silent mutations, recurrent CNAs and SVs that may be associated with either ABC or GCB COO are shown based on our validation cohort. An asterisk indicates significance at P<0.05. For genes with mutation hot spots or affected by CNVs or SVs, we considered these mutations separately from other missense variants. (B) Some genome-wide non-coding mutation peaks also showed cell-of-origin differences. Many of these are within genes that encode the immunoglobulin heavy and light chains. Unsurprisingly, the remaining genes overlap considerably with COO-associated genes that are also affected by coding mutations, mainly those affected by aSHM. The differential presence of aSHM activity, likely owing to expression differences, may explain why some of these genes are uniquely mutated in their respective subgroup. The *BCL2* locus had multiple peaks that were commonly mutated among GCB cases including multiple intronic regions that appear, based on H3K27Ac patterns, to coincide with an enhancer. These mutations were not restricted to cases with *BCL2* translocations. The *AICDA* locus, a novel aSHM target, is mutated mainly in ABC cases. The *BCL6* and *PAX5* super-enhancer was preferentially mutated in GCB cases. A peak in *GRHPR* near *PAX5* was more commonly mutated in ABC cases (Figure 3D). The *DNMT1* locus is near *S1PR2* and both of these peaks were enriched for mutations in GCB, indicating the potential for co-regulation of these genes using a common set of regulatory regions. (C) Genes with mutations significantly associated with one subgroup are shown above a heat map of the expression of several genes with strong COO-associated expression to highlight the mutual exclusivity between some gene pairs. In ABC, *NFKBIZ* and *MYD88* mutations were mutually exclusive relative to other mutations involved in NF-κB signalling. In GCB, *EZH2* and *MEF2B* hot spot mutations were common in *BCL2*-translocated cases and in those lacking mutations in *NFKBIA, MYC* and *SPEN*.

Within the 153 genomes, we identified between 1689 and 121,694 SSMs (median: 14,026). We searched genome-wide for patterns of mutation that may imply regulatory function without directly impacting protein sequence. To accomplish this, we implemented a new strategy to infer regions of arbitrary span with mutation density elevated above the local background. The method considers positions of pooled mutations from a cohort of cancer genomes excluding any variants within the CDS of genes (Methods). This analysis detected 4,386 peaks enriched for mutations that ranged from a single nucleotide to many kilobases (kb) in length (median length: 664 nucleotides). Using the pooled mutation set from all genomes, a randomly selected region should have, on average, 1.00 mutation per kb whereas these regions had a median mutation density of 10.3 per kb. Some hypermutated loci were represented by multiple peaks. For example, the *BCL6* locus and super-enhancer surrounding this region comprised 31 discrete peaks (Figure 3). Our analysis also revealed examples of non-coding loci with mutation peaks, for example the two adjacent long noncoding RNA (lncRNA) genes *NEAT1* and *MALAT1* and the miR-142 locus (Figure 3C). Mutations at each of these loci have been previously noted in other DLBCL and FL and their pattern is consistent with aSHM^27,28^.

**Figure 3:**
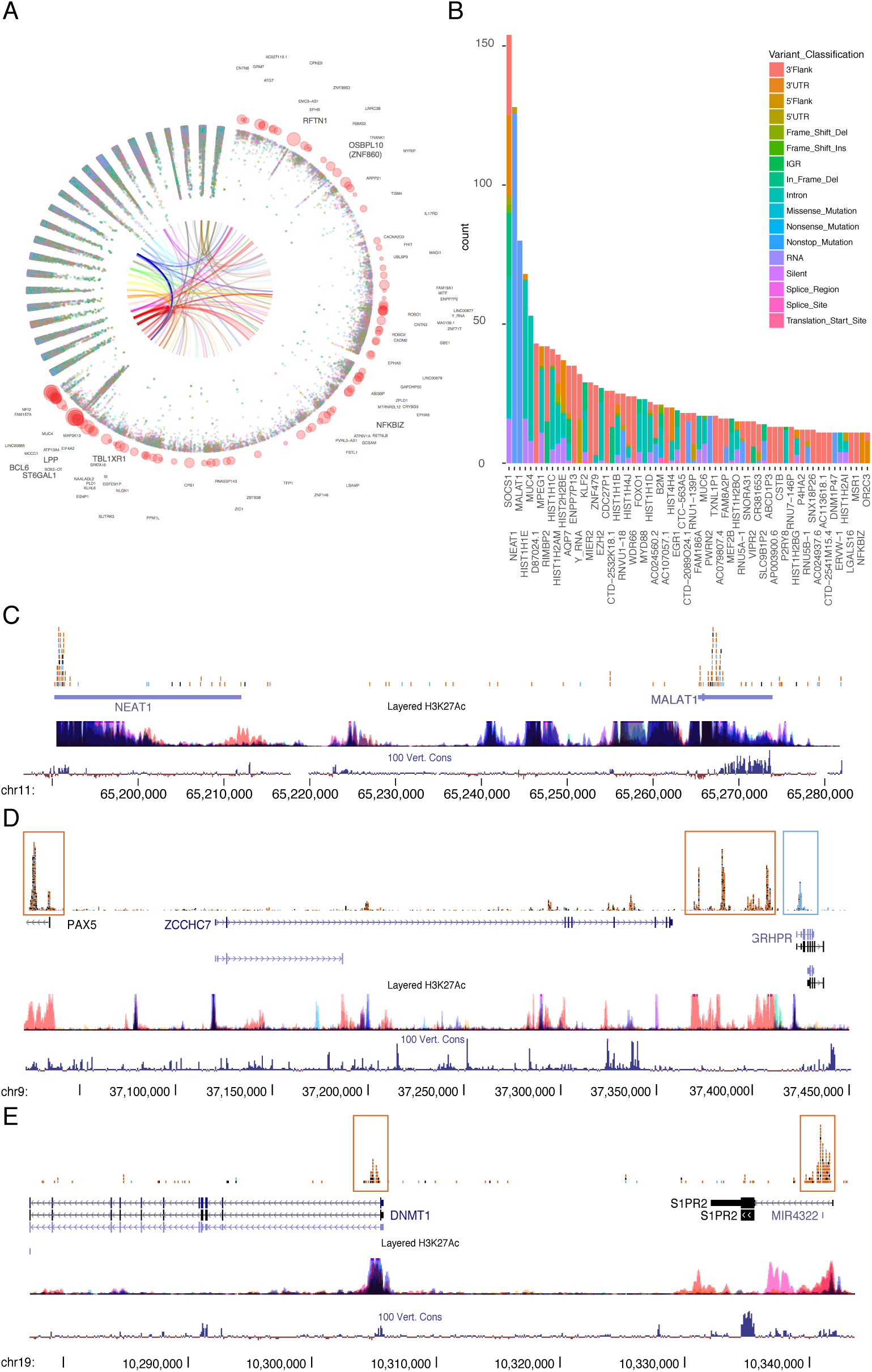
Annotation of novel recurrent mutations in non-coding and coding loci. (A) An overview of mutation peaks and the rainstorm representation of cohort-wide inter-mutation distance for chromosome 3. Red circles indicate mutation peaks identified by the wavelet approach. Internal arcs indicate SV breakpoints with distinct colours for each patient in the cohort. (B) Among the regions identified as enriched for SNVs through genome-wide analysis, the bulk of these affect intronic or intergenic regions or were near the transcription start site. Removing these may enrich for genes that are not merely affected by aSHM. Shown here are genes with one or more mutation peak and mutations in at least eight patients. The *NFKBIZ* locus had the strongest propensity for 3′UTR mutations among these remaining genes. This annotation-agnostic approach also detected sites in genes with mutation hot spots such as *EZH2, MYD88, B2M* and *FOXO1* and other genes known to act as drivers in DLBCL. (C) Multiple non-coding RNA genes are heavily mutated and show some focal enrichment, including the adjacent lincRNA genes *MALAT1* and *NEAT1.* (D) Similar to the *BCL6* super-enhancer, the locus containing *PAX5* and *ZCCHC7* harboured numerous mutation peaks. Peaks with mutations more common in GCB or ABC cases are boxed in orange and blue, respectively. (E) We found examples of mutation peaks proximal to genes that are commonly mutated in DLBCL such as *S1PR2.* In this example, *DNMT1* has a mutation pattern indicative of classical aSHM with most mutations directly downstream of the TSS.

To determine the suitability of our approach to identify mutation clusters relevant to DL-BCL, we extended our peak analysis to include all mutations including the coding region (CDS). We found a similar number of peaks (4,405), which comprised the bulk of the original non-coding peaks as well as peaks in genes with mutation hot spots such as *EZH2, FOXO1* and *MYD88* (Figure 3B). Aside from intergenic regions (2,244), the top three classes of annotation affected by peaks were 5′ flanking regions, 5′UTRs and introns. These are also the regions typically affected by aSHM and, as expected, virtually all of the known targets of this process were represented among these regions^12,14^. We also noted multiple genes with mutation patterns consistent with aSHM including the *AICDA* locus itself, *PRDM1, DNMT1,* and *ACTB* (Figure 3E and Extended Data Figure 3). If the mutations in these peaks were largely due to a single mutational process that preferentially acts in certain regions, we expected to find differences in the mutational signatures relative to the full set of SNVs. Interestingly, although the signature attributed to AID and POL*ν* activity was represented genome-wide and within the peaks, another signature that does not clearly correspond to any of the previously described signatures was unique to the peaks (Extended Data Figure 5).

To identify peaks with potential relevance in modulating transcription, we assessed the relationship between gene expression and the presence of mutations in nearby peaks. All genes with one or more proximal peak were tested for significant differences in expression between mutated and un-mutated cases (Extended Data Figure 6). Most of the protein-coding loci identified were known (including *SERPINA9, CD44, PIM1*) or the novel targets of aSHM we had identified (including *DNMT1* and *AICDA*). The correlation between expression and aSHM is typically attributed to an elevated AID activity at highly expressed genes and thus may act as a permanent marker of sustained expression of these genes rather than representing driver mutations. Regardless, the unprecedented breadth of mutations affecting potential regulatory regions including enhancers proximal to these genes suggests the possibility of a regulatory effect and this warrants further investigation. To enrich for genes with patterns unlikely to result from aSHM, we identified loci for which the most common variant annotation in each peak was not among the classes attributable to aSHM. Multiple genes showed distinct distributions of mutations seemingly inconsistent with aSHM. This could imply a different mutational process or the action of selective pressure to retain or alter function. Some of these genes had short 5′UTRs and thus had mutations within their CDS and even 3′UTRs (e.g. *MPEG1, HIST1H1C*). In longer genes, such as *NFKBIZ*, 3′UTR mutations cannot be readily attributed to aSHM and appear to indicate strong selective pressure.

### Recurrently Mutated Loci Associated With Cell-of-origin Subgroup

Our genome analysis uncovered a striking pattern of mutations in *NFKBIZ*, a gene that has been reported to act as an oncogene in DLBCL cases with copy number amplification affecting this locus^29^ though other somatic mutations affecting this region appear to be lacking. *NFKBIZ* was significantly more commonly mutated in ABC cases when the 3′UTR mutations are considered and even more strongly enriched in ABC when amplifications affecting this region are also considered (*P* = 2.15 × 10^-5^, Fisher’s Exact Test). Combining the genome data and results from targeted sequencing in a larger “validation” cohort, we confirmed our observation of a novel pattern of SSMs in the 3′UTR of *NFKBIZ* as well as some large indels and somatic structural variants (SVs) (Figure 5; Extended Data Figure 7). To demonstrate the improved resolution power of WGS to detect such mutations, we contrast these results to the large cohort of available exome data, which were uniformly re-analysed with the same methodology and show a much lower yield of these variants. We also compared the prevalence of *NFKBIZ* 3′UTR mutations in other lymphoid cancers with available WGS data including CLL, FL and BL. FL had the next highest prevalence with these mutations appearing in less than 3% of cases (Table 2). The number of cases also provided sufficient power to determine patterns of mutually exclusive genes within ABC and GCB. One of the few pairs of genes showing mutual exclusivity for mutation in ABC was *MYD88* and *NFKBIZ*, indicating a potentially redundant role of these two mutations in lymphomagenesis (Extended Data Figure 8).

**Figure 5:**
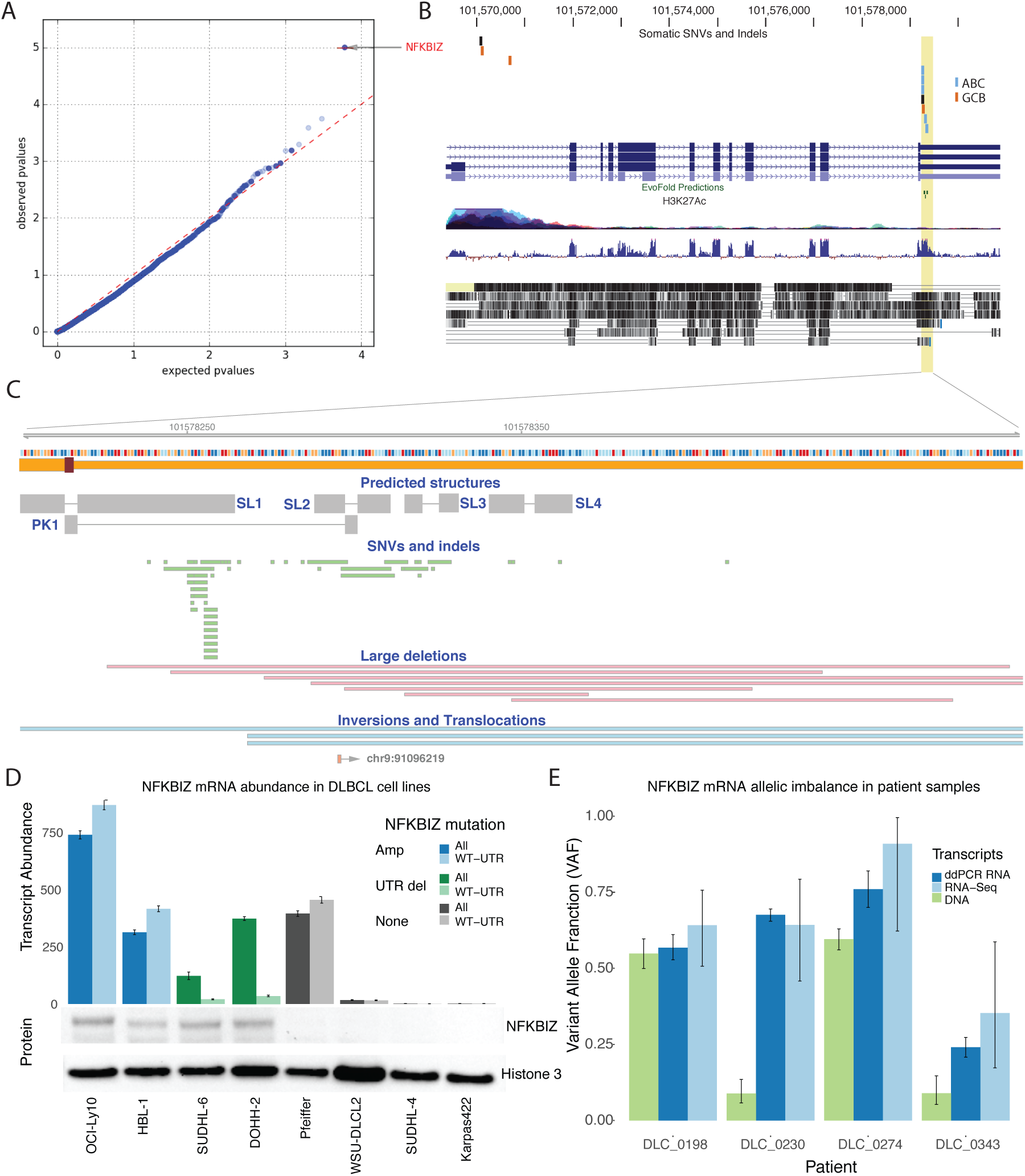
Mutations affecting the *NFKBIZ* locus and functional effects on mRNA and protein levels. (A) OncodriveFML^39^ quantile-quantile (QQ) plot comparing the expected and observed distribution of functional mutation (FM) bias P values for UTR mutations. *NFKBIZ* was the only gene with an FM bias q-value below 0.1. (B) These mutations clustered in a highly conserved region of the *NFKBIZ* 3′UTR and were significantly enriched in ABC cases (blue) relative to GCB cases (orange). (C) The mutated region of the 3′UTR showing the location of predicted structural features (grey) including stem-loops (SL) and a pseudoknot (PK). Individual mutations detected in the genomes or validation cohort are shown below. (D) A droplet digital PCR (ddPCR) assay was applied to eight DLBCL cell lines to determine *NFKBIZ* mRNA expression levels. Dark colours indicate total transcript counts and light colours indicate wild-type 3′UTR transcript counts. Cell line *NFKBIZ* mutations include amplifications (blue), 3′UTR deletions (green) or none (grey). One line (Pfeiffer) lacking any detectable *NFKBIZ* mutation had elevated *NFKBIZ* mRNA levels relative to un-mutated lines. We suspect this is due to a *STAT3* mutation, as previous studies suggest that *STAT3* plays a role in *NFKBIZ* activation^40,41^. Cell lines were also assessed by Western blot for IκBζ expression. Only mutant cell lines (green and blue) showed increased protein. (E) Comparison of variant allele fractions (VAFs) between DNA sequencing, RNA-Seq and ddPCR RNA assay for *NFKBIZ* mutant patients. RNA VAFs higher than the corresponding DNA VAFs indicate an allelic imbalance favouring the mutant allele.

**Table 2:** Prevalence of *NFKBIZ* 3′UTR mutations in DLBCL and other lymphoid cancers. Available WGS data from four lymphoid cancers was available from a combination of prior publications and unpublished *in house* data. Restricting to SSMs affecting the 3′UTR of *NFKBIZ*, the preva-lence was substantially higher in DLBCL than FL and appreciably lower in the DLBCL WES cohort. The prevalence in CLL and BL was below 1%.

| Disease | Data source | Total cases | % *NFKBIZ* 3'UTR mutated |
| --- | --- | --- | --- |
| DLBCL | Genome and Validation cohort | 449 | 9.13 |
| DLBCL | Published exome cohorts | 191 | 2.62 |
| FL | Published<sup>42</sup> and Unpublished, ICGC | 124 | 2.42 |
| BL | Unpublished and ICGC | 116 | 0.86 |
| CLL | Published<sup>15</sup> and unpublished | 144 | 0.69 |

The 3′UTR of *NFKBIZ* is highly conserved and the mutated region has been previously identified as a destabilising element that promotes rapid mRNA turnover^30^. We predict these mutations perturb this process thereby increasing mRNA longevity, which would in turn cause allelic imbalance in mutant cases. We constructed a structural model for the highly conserved region that contained the vast majority of SSMs (Extended Data Figure 9), which consists of one large stem-loop with several internal bulges and a smaller stem-loop with pairing in the loops forming a pseudoknot. Molecular dynamic simulations of selected mutations show that a subset appears to significantly change the structure relative to wild-type sequence (Extended Data Figure 9).

Higher mRNA levels were observed among the cases with 3′UTR mutations supporting their role in promoting mRNA abundance. To demonstrate the mutations promote elevated expression in *cis*, we searched for evidence of allelic imbalance. Of the cases with sufficient RNA-seq depth and at least one heterozygous SNP in *NFKBIZ*, we identified 25 cases with significant allelic imbalance. Roughly half of these cases had mutations that could impact either transcriptional or post-transcriptional regulation, with four containing a structural variation or indel affecting the 3'UTR and nine having one or more UTR SNV. This finding suggest that additional genetic or epigenetic alterations may also lead to allelic imbalance in *NFKBIZ* expression. We identified two DLBCL cell lines (DOHH-2 and SU-DHL-6) with *NFKBIZ* 3′UTR mutations and two lines (OCI-Ly10 and HBL-1) having *NFKBIZ* amplification.

To confirm this allelic imbalance, we implemented a ddPCR assay that separately quantifies mutant and wild-type *NFKBIZ* mRNAs. We tested mRNA extracted from eight DLBCL cell lines and a subset of the patient RNA samples and found that samples with *NFKBIZ* mutations or amplifications each had significantly higher mRNA levels. We confirmed in the two cell lines with *NFKBIZ* 3′UTR deletions (DOHH-2 and SU-DHL-6) that the mRNA represented a greater proportion of the mutant allele (Figure 5). When applying this assay to patient RNA samples, we compared the variant allele fractions (VAFs) for the mutation from our ddPCR assay and RNA-seq to the tumour DNA as determined by targeted capture sequencing (Extended Data Figure 10). The

RNA VAFs were higher than DNA in all *NFKBIZ* mutant patients, indicating increased expression of the mutant allele. Patient DLC_198 did not have a significant difference between RNA and DNA levels however, this can be attributed to a CNA of the *NFKBIZ* locus where an extra wild-type allele was present in this patient. By immunoblot, we confirmed high levels of IκBζ protein in these *NFKBIZ* mutant cell lines relative to lines lacking these events (Figure 5).

## Discussion

In this study, we have confirmed the recurrence and determined the specificity of non-silent mutations in a substantial number of genes to the molecular subgroups of *de novo* DLBCLs. Here, we focus our attention on some of the more common mutational features of DLBCL that were detectable only through the coverage of WGS data and integrative analysis with gene expression data from matched samples. Several recurrent sites of non-coding mutations were uncovered by a new algorithm. These include novel genes that are affected by aSHM and intergenic regions, particularly near active superenhancers, that appear to be mutated by the same process. The most striking finding within GCB DLBCL was the previously unappreciated recurrence of focal gains and complex events affecting the FC-**γ** receptor locus. The interplay between FC-**γ** receptors and cancer has general relevance because these proteins are directly involved in antibody-dependent cell-mediated toxicity (ADCC), which is triggered by monoclonal antibody-based (mAb) therapies including cetuximab, trastuzumab and rituximab^31^. Until recently, these have been studied in the context of Fc-**γ** receptor expression on effector cells and their interaction with mAbs on tumour cells in *trans.* In B-cell lymphomas, this situation is complicated by the presence of multiple Fc-**γ** receptor proteins on the malignant cells. Recent data have implicated common polymorphisms and gene expression differences in tumour tissue in variable response to rituximab but whether this was due to their effect on *cis* or *trans* interactions remained unclear. In CLL, *cis* interactions of Fc-**γ** receptor on malignant cells is associated with an elevated rate of internalisation of *FCGR2B* bound to IgG relative to its other family members^32^. Nonetheless, to date, recurrent somatic alterations promoting deregulated expression of FC-**γ** receptors have not been addressed.

We hypothesised that an imbalanced expression of FC-**γ** receptor proteins in malignant cells, due in part to the complex focal amplifications we have identified herein, attenuates the normal immune response to rituximab as has been seen with alternative isoforms and polymorphic variants of this gene. This was strongly supported by the significantly inferior outcome of *FCGR2B*-high GCB patients treated with R-CHOP (Figure 2) and is consistent with a smaller study that showed a correlation between FC**γ**RIIB protein staining and outcome in FL^33^. In light of this, alternative immunotherapy approaches may be warranted for this high-risk sub-population. Type II mAbs directed at CD20 or other proteins are not internalised by the same process and thus may be beneficial in these patients. Another potential avenue of exploration is direct targeting of FC**γ**RIIB alone or in combination^34^. Beyond somatic copy number alterations and possibly some influence from germline CNVs, we also identified an elevated level of SSMs in two introns of *FCGR2B* that may promote intron retention and lead to a truncated isoform. Further exploration of the role of genetic variation in producing *FCGR2B* over-expression or upsetting the balance of *FCGR2B* and *FCGR2C* in DLBCL is warranted.

In conjunction with *NFKBIZ* amplifications, which promote its expression, our data indicate an overall prevalence of 21.5% for mutations that may impact IκBζ protein abundance or function with 35 cases (10%) having at least one 3′UTR mutation. These were more common than coding changes or amplifications of this locus and strongly associate with the ABC subgroup whereas missense mutations were observed in both subgroups. Multiple studies have already attributed a 165 nt region in the UTR that harbours the bulk of the mutations we detected as destabilising elements^30,35^. The observation of allelic imbalance in many of the patient samples with 3′UTR mutations strongly implicates them in perturbing mRNA turnover but the functional mechanism is not clear. *NFKBIZ* is one of several genes subject to post-transcriptional regulation by the endoribonucleases Regnase-1 (Reg-1) and Roquin^35^. This process involves mRNA turnover and/or sequestration mediated by interactions between these proteins and specific stem-loops in the 3′UTRs of their targets^36^. *MYD88*, an adaptor protein that is commonly mutated in ABC DLBCLs, has been shown to be important for protecting *NFKBIZ* mRNA from this process^37^. Moreover, B-cell receptor signalling, which is active in most ABC DLBCLs, can also promote stabilisation of *NFKBIZ* mRNA via the UTR^30^.

The role of individual putative structural components in the 3′UTR of *NFKBIZ* in these individual processes has not been fully elucidated. The region we describe as containing a pseudoknot and comprising the first two stem-loops does not resemble the loop size and nucleotide properties that have been attributed to Roquin^38^, implicating instead the two distal elements in this interaction (SL3 and SL4). Reg-1 is encoded by *ZC3H12A,* one of the novel recurrently mutated genes identified through our meta-analysis (Extended Data Table 1). Confirming the mechanism whereby 3′UTR mutations impact the NF-κB pathway in DLBCL is highly relevant given the growing list of therapeutic strategies designed to inhibit this pathway directly or by perturbing upstream signalling events. Notwithstanding that mechanistic details remain unclear, we have demonstrated that higher mRNA and protein levels result from these mutations. To the best of our knowledge, recurrent 3′UTR mutations are the first example of a common somatic UTR alteration that can directly increase the expression of an oncogene.

## Acknowledgements

The British Columbia Cancer Agency Centre for Lymphoid Cancer gratefully acknowledges research funding support from the Terry Fox Research Institute, Genome Canada, Genome British Columbia, the Canadian Institutes for Health Research and the British Columbia Cancer Foundation. This work was supported by a Terry Fox New Investigator Award (1021) and an operating grant from the Canadian Institutes for Health Research (CIHR) to RDM, who is supported by a New Investigator award from CIHR and a Michael Smith Foundation for Health Research Scholar award. The genome sequence data and RNA-seq was produced with support from a contract with Genome Canada and Genome British Columbia led by JMC. MAM acknowledges the support of the Canada Research Chairs program and CIHR Foundation grant FDN-143288. The results published here are in whole or part based upon data generated by the TCGA Research Network: http://cancergenome.nih.gov/. We gratefully acknowledge TCGA and all providers of samples and resources for generating this valuable resource. The TCGA exome data was obtained through Database of Genotypes and Phenotypes (phs000178.v9.p8 and phs000450.v2.p1). We also acknowledge the ICGC MALY-DE project for providing access to their unpublished data. Aligned reads for those genomes were obtained through a DACO-approved project (to RDM) using a virtual instance on the Cancer Genome Collaboratory. Some of these data were produced as part of the Slim Initiative for Genomic Medicine (SIGMA), a joint US-Mexico project funded by the Carlos Slim Health Institute. AM is supported by fellowships from the Mildred-Scheel-Cancer-Foundation (German Cancer Aid), the MSFHR and Lymphoma Canada.The authors gratefully acknowledge the patient donors of samples used herein. We also thank Dr. Matthew Morin at Keyano College for the formal description of the rainstorm algorithm.

## Correspondence

Correspondence and requests for materials should be addressed to RDM.

